# Synaptopodin enables directional mechanoadaptation of integrin-based adhesions

**DOI:** 10.64898/2026.01.25.701551

**Authors:** Chengqing Qu, Yuxuan Huang, Shumeng Jiang, Xiangjun Peng, Yin-Yuan Huang, Pongpratch Puapatanakul, Zhiyang Xue, Kaye Brathwaite, Ewa Langner, Ethan Penn, Horacio D. Espinosa, Carmen M. Halabi, Elliot Elson, Jeffrey H. Miner, Guy M. Genin, Hani Y. Suleiman

## Abstract

The attachment of cells to their substrate through adhesion complexes is fundamental to tissue architecture and function. These adhesions are inherently optimized to resist shear forces parallel to the substrate, yet certain specialized cells must also withstand substantial perpendicular forces. How cells adapt their adhesion machinery to resist forces in different directions has remained unclear. In the kidney, podocytes experience perpendicular forces from pressurized filtrate flow while maintaining attachment to the glomerular basement membrane through integrin-based adhesions. Here we show that synaptopodin converts adhesions from shear-resistant to perpendicular force-resistant structures through coordinated reorganization of the actin cytoskeleton and adhesion complexes. Using an inertial force application system, we demonstrate that synaptopodin triggers force-dependent redistribution of β1-integrin to the cell periphery specifically in response to perpendicular loading, while synaptopodin-deficient cells lack this directional adaptation and detach. This mechanism operates in multiple cell types and is physiologically essential: synaptopodin-null mice subjected to elevated glomerular pressure develop significant proteinuria and podocyte foot process effacement. These findings reveal a molecular basis for directional mechanoadaptation, whereby a single protein enables cells to reconfigure their adhesion architecture in response to the direction of applied force.

## Introduction

The ability of cells to maintain specialized function in the presence of mechanical force is fundamental to multicellular life, but cells face very different mechanical challenges depending on their location and function in the organism. While most epithelial and endothelial cells are naturally equipped to resist shear forces that act parallel to their basement membrane, or roughly parallel to their stress fibers in the case of cells within tissues, some specialized cells must also withstand substantial forces pushing perpendicular to their surface. One site where this is particularly critical is the kidney glomerulus, in which specialized cells called podocytes, an essential component of the blood filtration barrier, are constantly resisting the perpendicular force of high-pressure ultrafiltration.^1–4^ Kidney disease from diverse causes, including genetic defect, injury, and toxins, can cause these cells to lose their grip on the glomerular basement membrane (GBM) and slough off into the urine, leading eventually to substantial pathology and in some cases kidney failure.^1,5^ Despite the importance of this mechanical adaptation, the ways that cells modify their internal architecture to resist these perpendicular forces is not known.

Synaptopodin is a key component and regulator of the podocyte cytoskeleton,^6–8^ with altered expression or stability serving as markers for podocyte related kidney disease.^9–11^ Varying forms of this actin-associated protein appear specifically in highly specialized cells that must maintain precise structural organization,^12^ including podocytes in the kidney and dendritic spine-containing neurons in the brain.^9^ While synaptopodin-deficient mice develop increased susceptibility to kidney injury and proteinuria under stress conditions, this cytoskeletal protein appears dispensable for normal podocyte development and function under baseline conditions.^10,11,13^ This makes synaptopodin particularly interesting as a potential mediator of cellular adaptation to mechanical stress, but we have an incomplete understanding of its precise molecular functions.

We hypothesized that synaptopodin serves as a mechanosensitive regulator of adhesion complexes, enabling podocytes to resist perpendicular forces from the pressurized flow of filtrate. These integrin-based adhesions, termed focal adhesions in culture although likely organized differently *in* vivo, naturally anchor cells against shear forces parallel to the substrate.^14^ However, podocytes face the unusual challenge of withstanding substantial perpendicular forces from fluid filtrate that could detach them from the GBM. Our model proposes that synaptopodin strengthens adhesion points through two coordinated mechanisms: reinforcing the actin bundles that connect to adhesions and modulating the distribution of adhesion complexes in response to mechanical stress.

## Results

### Synaptopodin Enables Podocytes to Resist Perpendicular Forces

Podocytes face a unique mechanical challenge: they must maintain stable adhesion to the GBM while resisting substantial perpendicular forces from blood pressure driving the flow of ultrafiltrate through their filtration slit diaphragms. To investigate how podocytes accomplish this, we developed two complementary experimental systems that isolate the key mechanical forces these cells experience *in vivo*. For perpendicular forces, we cultured primary podocytes on laminin-521-coated polyacrylamide hydrogels^15^ and applied calibrated centrifugal forces using a temperature-controlled swinging bucket centrifuge. To examine lateral shear forces separately, we designed a microfluidic chamber that generates controlled laminar flow across the cell surface (Fig. 1a and Supplementary Fig. 1a).

**Fig. 1.**
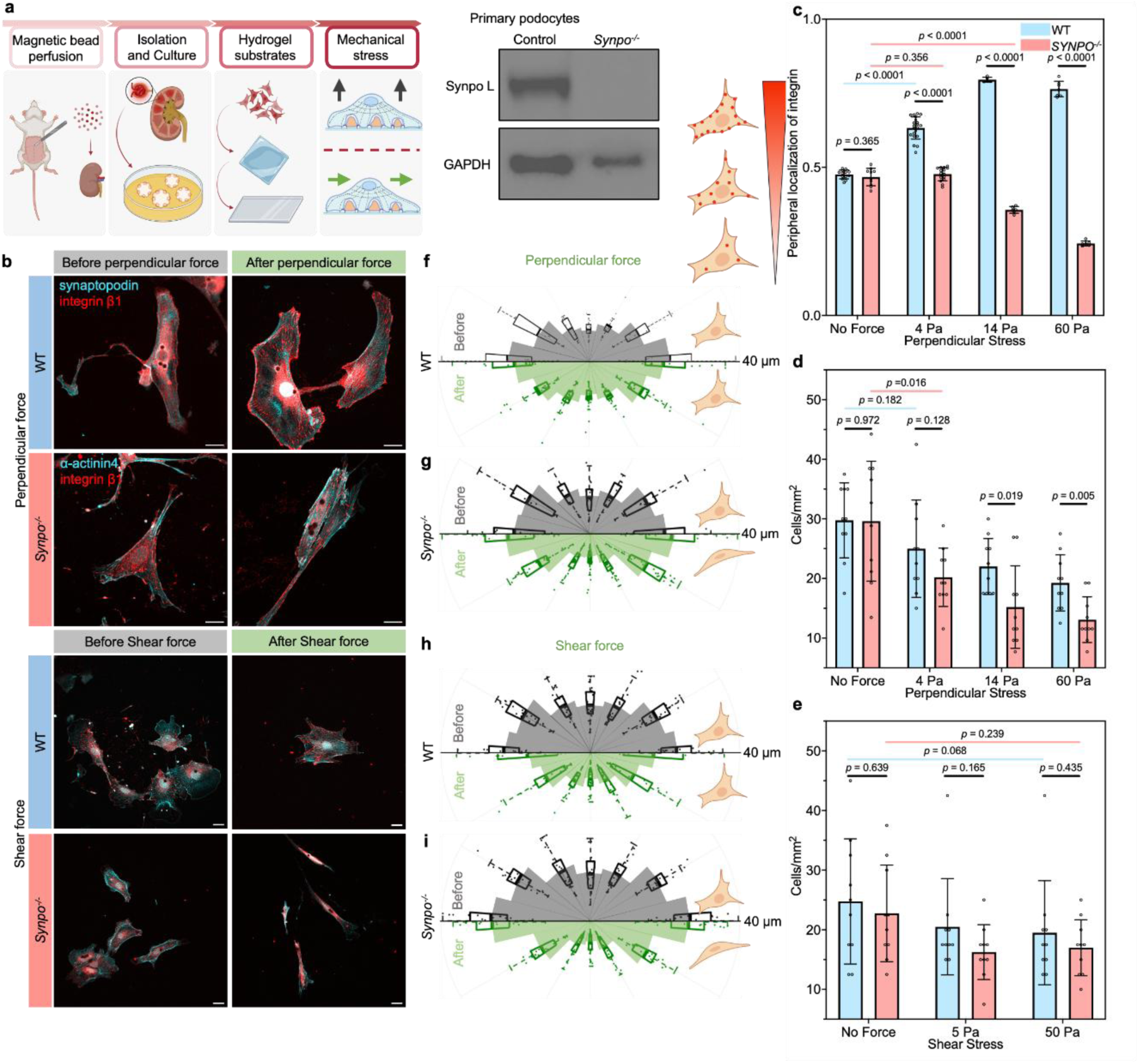
Synaptopodin enables podocytes to resist perpendicular forces through directional reorganization of focal adhesions. (**a**) Experimental approach: primary podocytes cultured on laminin-521-coated hydrogels were subjected to perpendicular forces via centrifugation or lateral shear forces via microfluidic flow. Inset: Western blot confirming synaptopodin expression in WT but not *Synpo*^⁻/⁻^ podocytes. (**b**) Representative immunofluorescence images of WT and *Synpo*^⁻/⁻^ podocytes stained for synaptopodin (green) and β1-integrin (magenta) under baseline, perpendicular force, and lateral shear force conditions. WT cells redistribute β1-integrin to the cell periphery specifically in response to perpendicular loading; *Synpo*^⁻/⁻^ cells fail to reorganize adhesions under either loading condition. Scale bars: 20 µm. (**c**) Quantification of focal adhesion distribution (peripheral localization index) as a function of perpendicular stress magnitude. WT cells show progressive peripheral redistribution up to 14 Pa; organization is disrupted at 60 Pa, indicating an upper mechanical limit. *Synpo*^⁻/⁻^ cells show no systematic reorganization at any force level. (**d**) Cell retention under increasing perpendicular force. WT podocytes maintain strong adhesion, whereas *Synpo*^⁻/⁻^ cells detach readily. (**e**) Cell retention under lateral shear force. Neither genotype shows significant detachment, indicating that synaptopodin specifically mediates perpendicular force resistance. (**f–i**) Rose diagrams depicting cell shape distributions. WT cells maintain stable aspect ratios under both perpendicular (**f**) and lateral shear (**g**) loading. *Synpo*^⁻/⁻^ cells undergo marked elongation under both perpendicular (**h**) and lateral shear (**i**) forces, indicating a general cytoskeletal defect that translates to adhesion failure only under perpendicular loading. Data represent mean ± SEM; *n* ≥ 15 independent experiments per condition. Scale bars: 20 µm.

The presence of synaptopodin, confirmed by Western blot analysis (Fig. 1a), proved critical for cells to maintain attachment to their substrate under perpendicular loading. When subjected to increasing perpendicular forces, primary podocytes isolated from wild-type (WT) mice showed strong resilience, with about 83% of cells remaining attached even under 14 Pa stress; these cells expressed the long isoform of synaptopodin. In contrast, primary podocytes from synaptopodin knockout (*Synpo^-/-^*) mice,^11,13^ with neither the long nor short isoform of synaptopodin, showed significantly greater detachment, with only 33% of cells remaining attached under the same conditions (*p* < 0.001; Fig. 1d). This difference was specific to perpendicular forces: neither WT nor *Synpo^-/-^* podocytes showed significant detachment under lateral shear flow, suggesting synaptopodin specifically enables resistance to perpendicular forces (Fig. 1e).

### Synaptopodin Mediates Force-Dependent Reorganization of Focal Adhesions

To understand how synaptopodin enables this mechanical adaptation, we examined the distribution of integrin-based focal adhesions (FAs) under force. WT primary podocytes responded to perpendicular force by dramatically redistributing β1 integrin to their periphery, whereas *Synpo*^-/-^ cells maintained the baseline uniform distribution of FAs (Fig. 1b). This redistribution appeared force-dependent, with WT cells showing progressive FA reorganization with increasing stress (from 0-14 Pa), resulting in increasingly peripheral distribution as measured by our FA localization index (Fig.1c). However, high stress (60 Pa) disrupted this organization, suggesting a mechanical limit to this adaptation. In contrast, *Synpo*^-/-^ cells showed no systematic reorganization of FAs at any force level, indicating that synaptopodin is essential for this mechanoadaptive response (Fig.1b,c).

### Synaptopodin Maintains Cell Shape and Adhesion Under Mechanical Load

Since mechanical forces can significantly alter cellular architecture, we investigated how synaptopodin affects cell shape stability of primary podocytes under load. WT cells maintained a consistent aspect ratio (length/width ratio of 2-3) under both baseline conditions and mechanical loading, becoming only slightly more rounded under perpendicular force and slightly more polarized under lateral shear (Fig. 1f,h). In contrast, *Synpo^-/-^* cells underwent substantial elongation under both loading conditions, with their aspect ratio increasing from 2-3 at baseline to 4-5 under load (Fig. 1g,i). This indicates that synaptopodin is required for maintaining cell shape stability under mechanical stress generally.

The adhesion phenotype was direction-specific. While *Synpo^-/-^* cells showed shape distortion under both perpendicular and lateral shear loading, they detached preferentially under perpendicular force (Fig. 1d,e). WT cells remained well-attached under both conditions. This dissociation between shape and adhesion phenotypes suggests that synaptopodin’s most critical function is not simply maintaining cell shape, but rather enabling adhesion complexes to resist perpendicular detachment forces. The shape instability in *Synpo^-/-^* cells may reflect a general cytoskeletal defect, but this only translates to adhesion failure when cells face perpendicular loading.

### Synaptopodin Enables U2OS Cells to Resist Perpendicular Forces

To test whether synaptopodin’s role in resisting perpendicular forces extends beyond specialized kidney cells, we investigated cells of the U2OS human osteosarcoma cell line,^16^ which we discovered express synaptopodin despite having no obvious physiological need to resist perpendicular forces. Western blot analysis confirmed that U2OS cells express both long and short synaptopodin isoforms (Fig. 2a). Using CRISPR-Cas9, we generated *SYNPO* knockout U2OS cells and confirmed complete loss of both isoforms (Fig. 2a). Additionally, *SYNPO^-/-^* U2OS cells showed reduced levels of α-actinin-4 (Fig. 2a), similar to cultured *Synpo^-/-^*podocytes.^10,11^

**Fig. 2.**
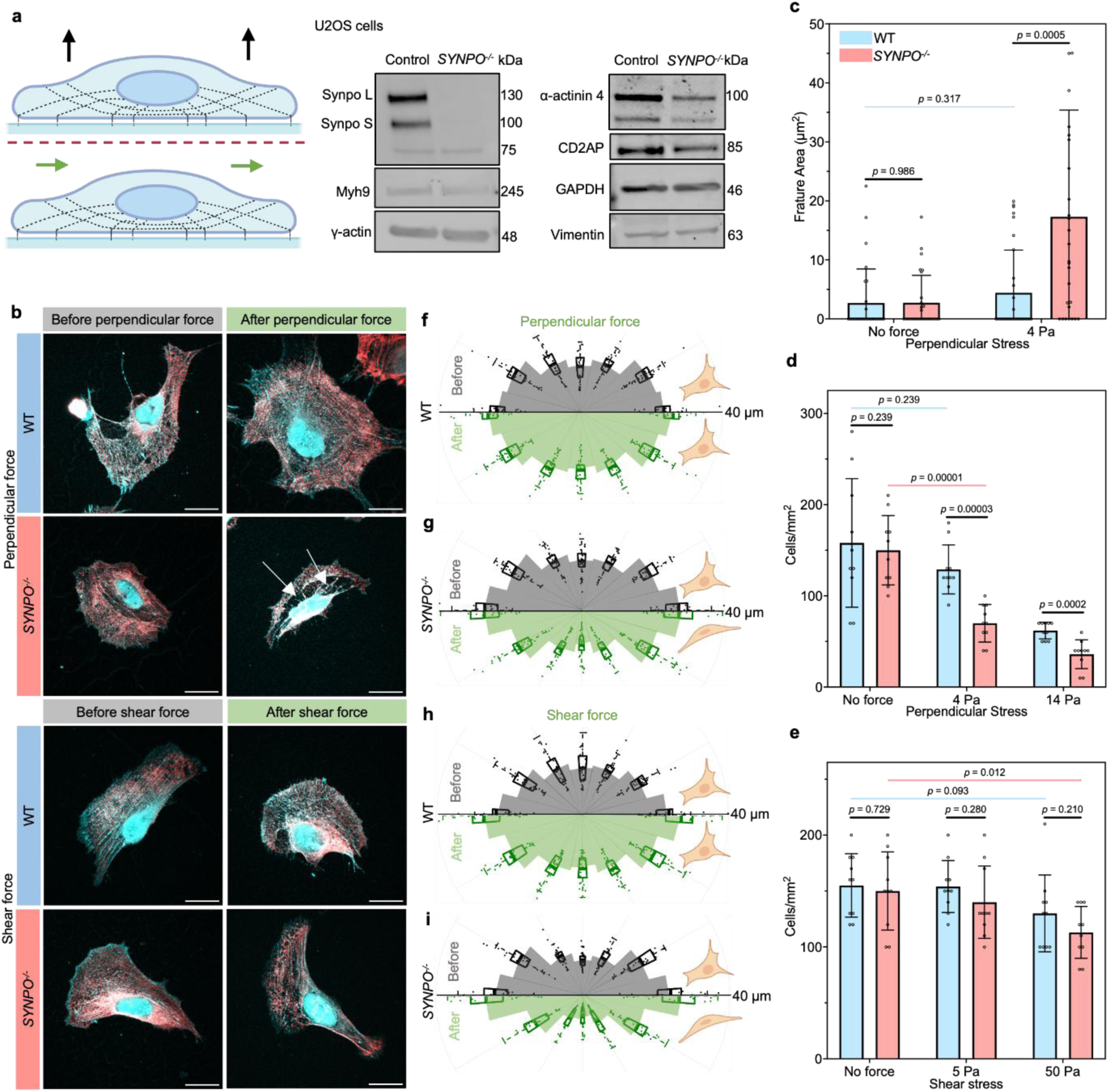
Synaptopodin-dependent perpendicular force resistance is conserved in U2OS cells and requires cytoskeletal integrity. **(a)** Force application conditions. Western blots demonstrate that WT U2OS cells express both long (SYNPO-L) and short (SYNPO-S) synaptopodin isoforms, while CRISPR-generated *SYNPO*^⁻/⁻^ cells lack both. α-Actinin-4 (ACTN4) levels are reduced in *SYNPO*^⁻/⁻^ cells, consistent with synaptopodin’s role in stabilizing this binding partner. **(b)** Representative immunofluorescence images of WT and *SYNPO*^⁻/⁻^ U2OS cells stained for non-muscle myosin IIA (MyoIIA; red) and ACTN4 (cyan) before and after perpendicular force application. WT cells maintain organized sarcomere-like structures (SLSs) containing both proteins under all conditions. *SYNPO*^⁻/⁻^ cells display small perinuclear gaps in the cytoskeletal network at baseline; these gaps expand into severe cytoskeletal fractures under perpendicular loading, while intact regions retain SLS organization. **(c)** Quantification of fractured cytoskeletal area per cell. *SYNPO*^⁻/⁻^cells exhibit significantly larger fracture zones following perpendicular force application compared to WT cells. **(d)** Cell retention under perpendicular force. At 4 Pa, WT cells remain well-attached, whereas *SYNPO*^⁻/⁻^ cells show substantial detachment (*p* < 0.001). Both genotypes detach extensively at 14 Pa, indicating a lower force threshold in U2OS cells compared to podocytes. **(e)** Cell retention under lateral shear force shows no significant difference between WT and *SYNPO*⁻/⁻ cells (*p* > 0.05), confirming that synaptopodin specifically mediates perpendicular force resistance. **(f, g)** Rose diagrams depicting WT cell shape under perpendicular **(f)** and lateral shear **(g)** forces. WT cells maintain stable aspect ratios (∼2) under both loading conditions. **(h, i)** Rose diagrams of *SYNPO*^⁻/⁻^ cell shape under perpendicular **(h)** and lateral shear **(i)** forces reveal that mutant cells elongate substantially, mirroring the phenotype observed in *Synpo*^⁻/⁻^ podocytes. Data represent mean ± SEM; *n* ≥ 15 independent experiments per condition. Scale bars: 20 µm.

When subjected to perpendicular forces, *SYNPO^-/-^* U2OS cells showed similar defects to *Synpo^-/-^* podocytes. Under 4 Pa stress, while WT U2OS cells maintained their attachment with only 7% cell loss, *SYNPO^-/-^* cells showed significant detachment with nearly 47% of cells lost (*p* < 0.001, Fig. 2d). At higher stress (14 Pa), both WT and mutant cells showed extensive detachment, indicating that U2OS cells have a lower force threshold than podocytes. Under lateral shear forces, WT U2OS cells remained well-attached (86%), while *SYNPO^-/-^*cells showed modest detachment (80%, *p > 0.05)* at 50 Pa (Fig. 2e).

The mechanical vulnerability of *SYNPO^-/-^* U2OS cells was manifested in both cytoskeletal integrity and cell shape stability. Even before force application, *SYNPO^-/-^* cells showed gaps between the nucleus and cell periphery, visualized by non-muscle myosin IIA (MyoIIA) and α-actinin-4 (ACTN4) staining. These gaps developed into severe cytoskeletal fractures when perpendicular force was applied (Fig. 2b,c) and were evident after force application (Supplemental Fig.1 b,c). Loss of cytoskeletal integrity was accompanied by cell shape distortion. Whereas WT U2OS cells maintained a consistent aspect ratio of around 2 under perpendicular force (Fig. 2f,h), *SYNPO^-/-^* cells elongated significantly, with their aspect ratio increasing from 2.5 at baseline to approximately 4 under load (Fig. 2g,i).

These findings, parallel to those in podocytes, suggest that synaptopodin’s ability to maintain cellular architecture under perpendicular force is a fundamental mechanical adaptation that functions even in cells that do not typically experience such forces. The consistent pattern of cytoskeletal disruption and shape distortion in diverse cells lacking synaptopodin points to a conserved mechanism.

### Lack of Synaptopodin Impairs U2OS Cell Motility

To test the degree to which synaptopodin affects adhesion-associated cell function, we applied a cell migration assay that monitored closure of a gap between populations of either WT or *SYNPO*^-/-^ U2OS cells. The gap between WT U2OS colonies narrowed visibly within 9 hours and disappeared within 24 (Supplemental Fig. 1d,e), but the gap between *SYNPO*^-/-^ U2OS cells had not begun to narrow at 9 hours and was still evident at 24 hours. This reinforces that synaptopodin contributes to adhesion related cell function.

### Increased Blood Pressure Reveals Mechanical Vulnerability in *Synpo^-/-^* Mice

To test the potential physiological relevance of synaptopodin’s role in resisting mechanical forces, we examined how *Synpo*^-/-^ mice respond to increased blood pressure, which elevates perpendicular forces on podocytes due to increased pressurized flow of filtrate. We induced hypertension using an adeno-associated virus serotype 8 (AAV8) that drives renin expression, with an AAV8 expressing Green Fluorescent Protein (GFP) serving as a control. Both AAVs showed comparable expression in the livers of WT and *Synpo*^-/-^ mice (Fig. 3a), confirming successful transduction.

**Fig. 3.**
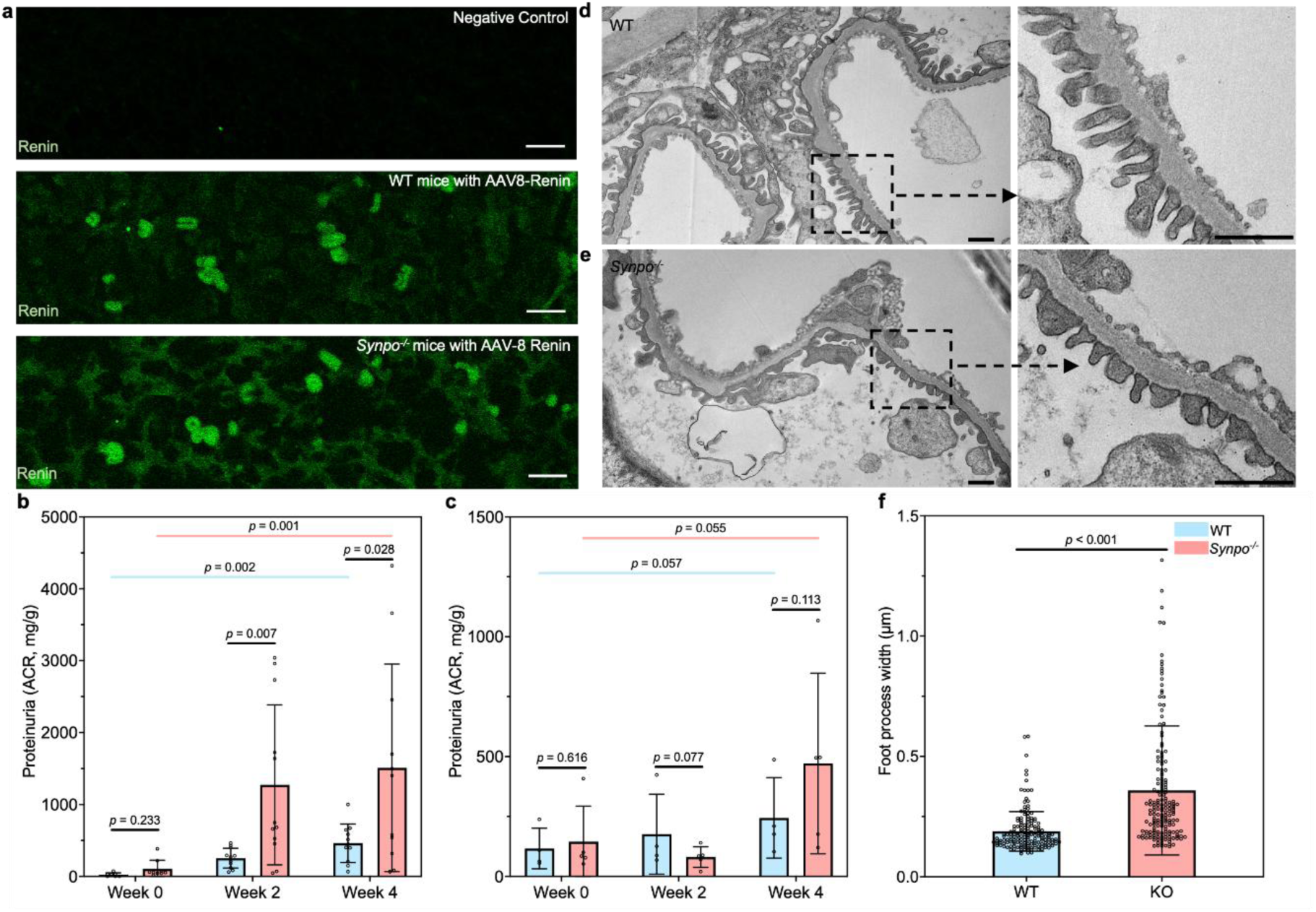
Synaptopodin is essential for maintaining podocyte architecture under elevated glomerular pressure *in vivo*. **(a)** Validation of hypertension model. Representative immunostaining for renin in liver cryosections from uninjected WT mice, AAV8-Renin-injected WT mice, and AAV8-Renin-injected *Synpo^-/-^* mice. Robust renin expression is detected in hepatocytes of both injected genotypes, confirming successful viral transduction and ectopic renin production. **(b)** Albumin-to-creatinine ratios measured at 0, 2, 4, and 6 weeks following AAV8-Renin injection. *Synpo^-/-^*mice develop significant proteinuria beginning at week 4 (*p* < 0.05 vs WT), whereas WT mice maintain normal kidney function despite equivalent mean arterial blood pressure elevation (15-20 mm Hg increase, to 90-100 mm Hg). **(c)** Control AAV8-GFP injections do not induce proteinuria in either genotype, confirming that filtration barrier dysfunction in *Synpo^-/-^* mice results from elevated pressure rather than viral infection itself. **(d, e)** Representative transmission electron micrographs of glomerular capillary loops from AAV8-Renin-injected WT **(d)** and *Synpo^-/-^* **(e)** mice. WT podocytes maintain normal foot process architecture with regularly spaced interdigitations. *Synpo^-/-^* podocytes display widened foot processes and extensive effacement, indicating loss of normal cytoskeletal organization under mechanical stress. **(f)** Quantification of foot process width. *Synpo^-/-^* mice exhibit significantly increased foot process width compared to WT controls following AAV8-Renin treatment, consistent with cytoskeletal dysfunction under elevated perpendicular filtration forces. These *in vivo* findings demonstrate that synaptopodin-mediated perpendicular force resistance observed *in vitro* is physiologically essential for podocyte survival under hemodynamic stress. Data represent mean ± SEM; n ≥ 4 mice per group. Scale bars: (a) 20 µm (d,e) 1 µm.

Mean arterial blood pressure (MAP) measurements showed an increase of 15 to 20 mm Hg in AAV8-Renin-infected mice, which had an average MAP of 92.3 mm Hg vs. an average MAP of 78.7 mm Hg in AAV8-GFP-infected mice. The response to increased blood pressure differed in WT vs. *Synpo*^-/-^ mice. Whereas WT mice maintained normal kidney function, *Synpo*^-/-^ mice developed significant proteinuria under elevated blood pressure, with albumin-to-creatinine ratios increasing markedly from week 4 after AAV8-Renin administration (*p* < 0.05, Fig. 3b). *Synpo*^-/-^ mice receiving control AAV8-GFP maintained normal kidney function (*p* > 0.05, Fig. 3c). Because proteinuria can be caused by podocyte loss, we quantified podocyte number relative to glomerular area. After AAV-Renin administration, *Synpo*^-/-^ mice showed significantly fewer podocytes than WT mice (Supplemental Fig. 2a-c). Electron microscopy confirmed that *Synpo*^-/-^mice subjected to increased blood pressure showed loss of normal foot process architecture, with average foot process width significantly greater than controls (Fig. 3d-f). Defects in adhesion and structural deterioration under increased mechanical load indicate that synaptopodin is essential for maintaining podocyte architecture against elevated mechanical forces *in vivo*.

### Synaptopodin Forms Mechanically Oriented Structures with Actin and Myosin

To understand the structural basis for synaptopodin’s mechanical role, we examined its cytoskeleton organization using high-resolution microscopy. In podocyte foot processes *in vivo*, we discovered that synaptopodin forms distinctive structures wrapping around actin filaments that extend from the central actin bundle to the basal surface where the cell attaches to the GBM (Fig. 4a), consistent with our earlier focused ion beam scanning electron microscopy (FIB-SEM) imaging of the podocyte actin network.^17^ In cultured primary podocytes, synaptopodin and MyoIIA formed alternating patterns reminiscent of sarcomeres (Fig. 4b), particularly at the cell periphery where shear lag effects amplify mechanical stresses.^15^ These sarcomere-like structures oriented toward sites of cell adhesion, with β1 integrin clustering at their termini.

**Fig. 4.**
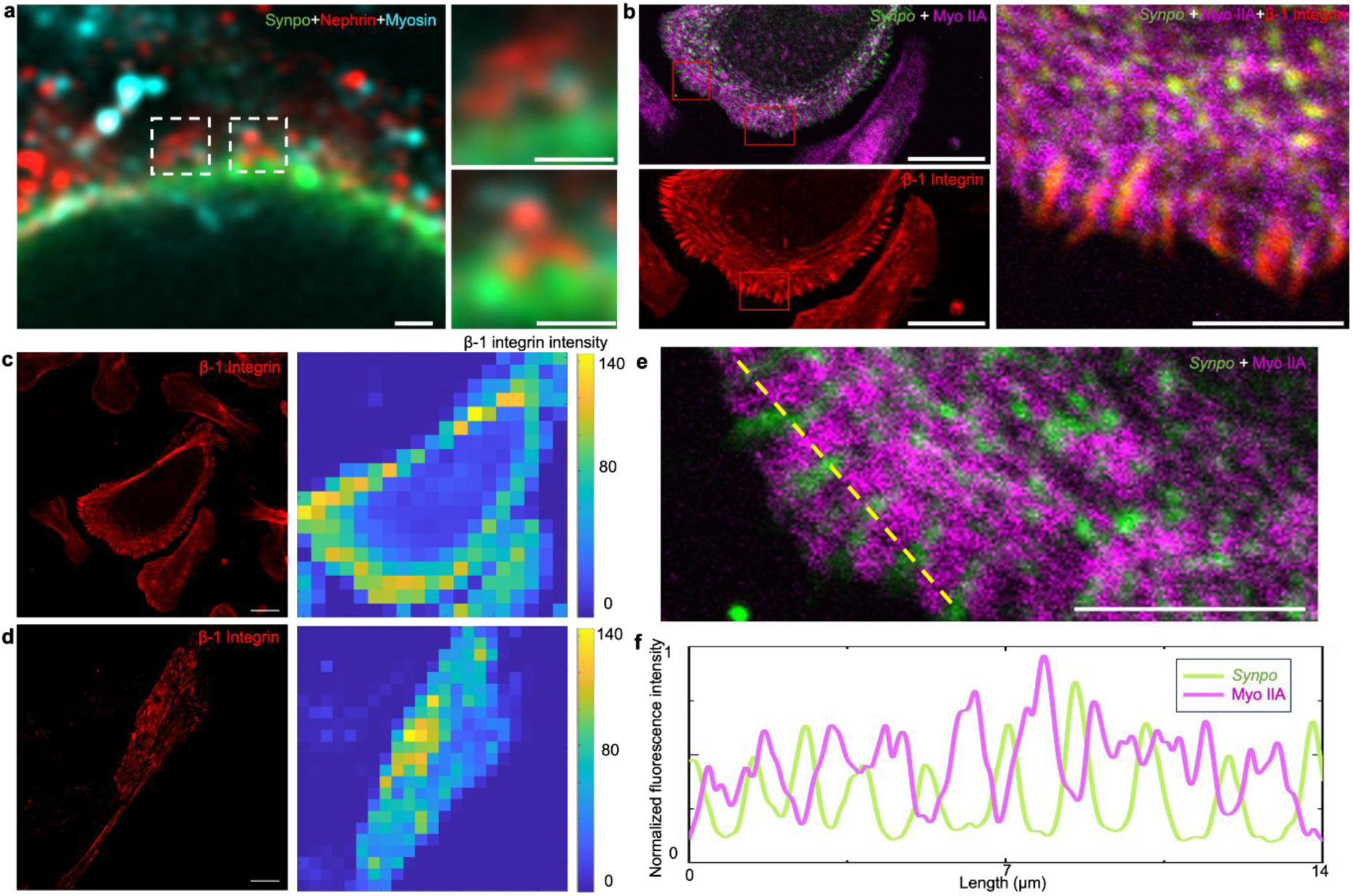
Synaptopodin organizes force-transmitting structures at adhesion sites, and a computational model recapitulates synaptopodin-dependent mechanoadaptation. **(a)** Expansion microscopy of WT mouse kidney cortex immunostained for synaptopodin (red), nephrin (green), and MyoIIA (cyan). Low-magnification image (scale bar: 0.2 µm) shows glomerular architecture; boxed regions are magnified at right (scale bar: 0.2 µm). Synaptopodin forms distinctive triangular wrapping structures around actin filaments that extend from central actin bundles to the basal surface where foot processes contact the glomerular basement membrane, consistent with a role in transmitting forces to adhesion sites. (b) Immunofluorescence of a WT primary podocyte after perpendicular force application, stained for synaptopodin (green), MyoIIA (purple), and β1-integrin (separate panel). Synaptopodin and MyoIIA form alternating sarcomere-like structures (SLSs) that are maintained at the cell periphery under load. β1-integrin accumulates at the periphery, colocalizing with synaptopodin at the termini of actin cables but remaining spatially distinct from MyoIIA. This molecular organization suggests a division of labor in which actomyosin generates contractile force while synaptopodin-organized structures transmit force to adhesion complexes. Scale bar: **(b)** 20 µm, ROI 5 µm. **(c)** Immunofluorescence and corresponding heat map of a WT primary cell after perpendicular force application stained for β1-integrin. β1-integrin formed clusters at the periphery. (d) Immunofluorescence and heatmap of a *Synpo^-/-^* primary cell after perpendicular force application, stained for β1-integrin. β1-integrin was not peripherally distributed. (e) The left ROI in **(b)** with an added dashed 14 µm line used to plot synaptopodin (green) and MyoIIA (purple). **(f)** Normalized fluorescence intensity plot of synaptopodin (green) and MyoIIA (purple) signal along the 14 µm line in **(e)**, showing a well-organized SLS.

Synaptopodin and β1 integrin colocalized at these sites, whereas MyoIIA remained distinctly separate (Fig. 4a,b). Compared with WT primary podocytes, *Synpo^-/-^* primary podocytes did not exhibit β1 integrin clustering at the periphery under perpendicular force. (Fig. 4c,d). The sarcomere like structures in WT primary podocytes have both synaptopodin and MyoIIA, but in an alternating, nonoverlapping pattern (Fig. 4e,f). This molecular organization suggests a model in which synaptopodin helps organize force-transmitting structures that connect the actomyosin machinery to adhesion sites at the cell periphery.

To understand synaptopodin regulation of myosin IIA, we examined myosin IIA reorganization following transfection of the long isoform of synaptopodin in cultured *Synpo^-/-^* primary podocytes (Supplemental Fig. 3a-b). To achieve single-cell resolution while maintaining high viability in these sensitive primary cells, we employed nanofountain probe electroporation (NFP-E), a localized electroporation technique that delivers cargo into individual cells with minimal perturbation to neighboring cells.^18,19^ This approach generates mosaic expression patterns, enabling direct comparison of transfected and non-transfected cells within the same culture conditions, providing internal controls that eliminate confounding factors stemming from batch-to-batch variability in primary cell preparations. NFP-E proved particularly advantageous for primary podocytes, which are difficult to transfect using conventional methods and exhibit poor survival following bulk electroporation.

In the absence of synaptopodin transfection, myosin IIA presented no clear organization (Supplemental Fig. 3c,d). Following single-cell transfection of neighboring cells, myosin IIA adopted a sarcomeric pattern, alternating with synaptopodin (Supplemental Fig. 3e,f), indicating that synaptopodin enables organization of contractile machinery in primary podocytes.

### Results Are Consistent with a Model of Synaptopodin Participating in Lateral Shear-Strengthening of Adhesion Complexes

To develop a mechanistic understanding of how synaptopodin enables perpendicular force resistance, we constructed a computational model of cell detachment under inertial loading. The model represents the cell as a mechanical system in which discrete stress fibers, modeled as elastic cables, connect a rigid nucleus to focal adhesions at the cell periphery (see Methods). Each focal adhesion was governed by force-dependent catch-slip kinetics, with bond dissociation rates modulated by both normal (i.e., perpendicular) and lateral shear force components (Fig. 5a).

**Figure 5.**
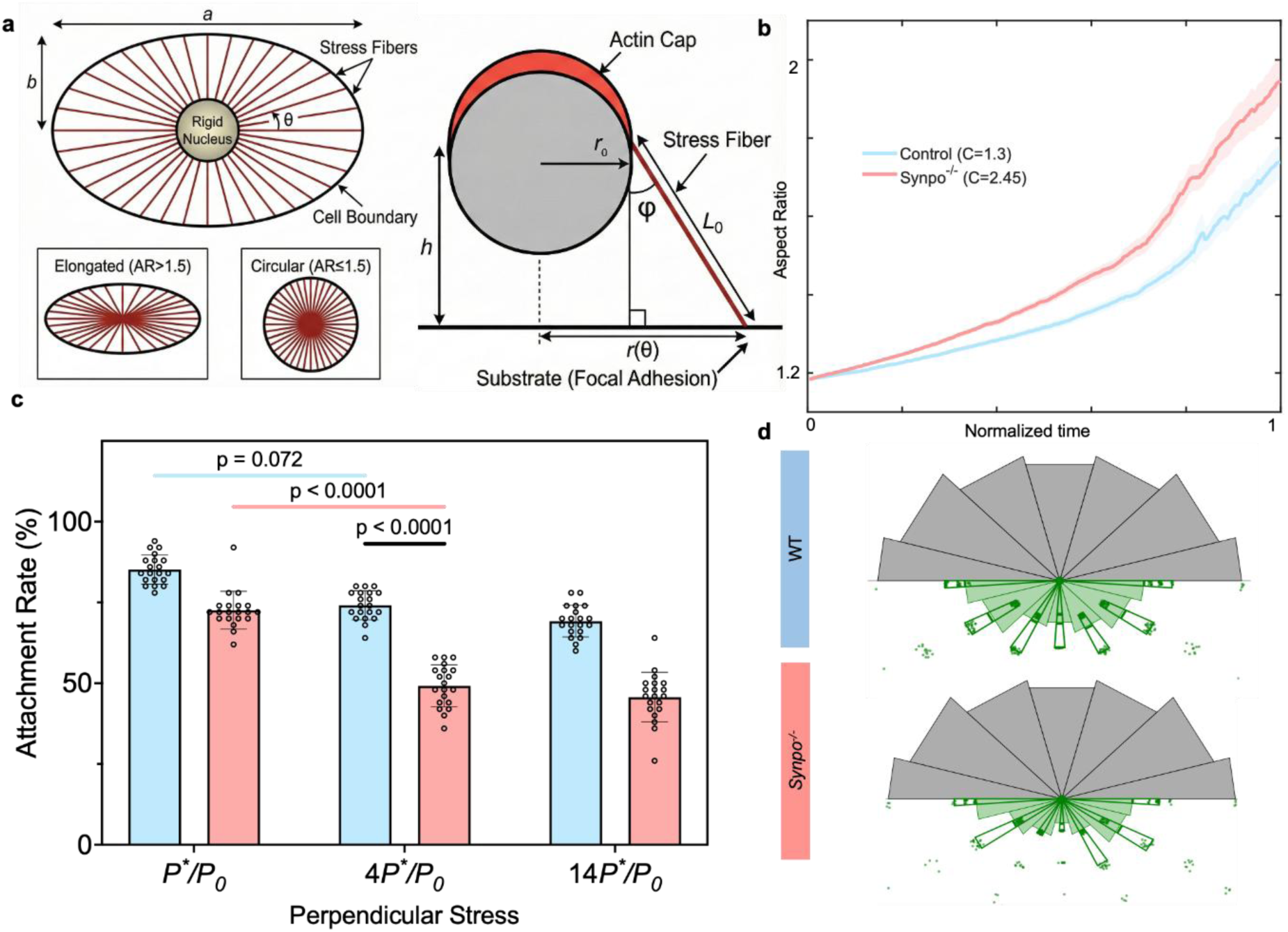
Computational modeling of cell detachment dynamics and stress fiber architecture. **(a)** Schematic of the geometric model setup. The cell boundary is defined as an ellipse with major axis *α* and minor axis *b*. The top-down view (left) illustrates stress fiber distribution, which is weighted for elongated cells (Aspect Ratio, *AR* > 1.5) and uniform for circular cells (*AR* ≤ 1.5). The cross-sectional view (right) depicts the rigid nucleus with an actin cap of radius *r*_0_ at height ℎ, connected to the substrate via stress fibers of length *L*_0_ at angle *ϕ*. Simulation results show the evolution of the cell Aspect Ratio over normalized time during perpendicular force loading. The plot compares adhesion strength conditions for Control (*C* = 1.3, blue) and Synpo^-/-^ (*C* = 2.45, red), where *C* modulates the molecular bond parameter. **(c)** Quantification of the cell attachment rate (percentage of cells remaining attached) under varying magnitudes of perpendicular stress (*P*^∗^/*P*_0_). The bar graph compares Control and *Synpo^-/-^* cells across increasing force loads (1 × to 14 × base stress). Statistical significance is indicated by *P*-values. **(d)** Rose diagrams demonstrate the angular distribution of attached stress fibers for WT and *Synpo^-/-^* cells before and after the loading. Each radial block represents the quantity of bonded fibers remaining at specific orientations relative to the cell axis.

A central feature of the model is a parameter *C* that represents the degree to which normal forces transmitted through stress fibers are converted to lateral shear forces at focal adhesion sites. This conversion depends critically on the mechanical integrity of actin cables: when cables are reinforced, they resist lateral displacement under perpendicular loading, distributing forces across multiple adhesion sites and converting a portion of the normal load into lateral shear stress that stabilizes catch bonds. We hypothesized that synaptopodin provides this reinforcement by bundling actin filaments and organizing them into mechanically coherent structures.

We fit the model to experimentally measured cell detachment rates under increasing perpendicular force using *C* as the sole free parameter. With *C* = 1.3 for WT cells and *C* = 2.45 for synaptopodin-deficient cells, the model accurately recapitulated the experimental detachment curves (Fig. 5c). The higher *C* value in WT cells reflects a test of the hypothesis of enhanced force conversion: synaptopodin-reinforced actin cables transmit perpendicular loads to focal adhesions in a manner that engages catch-bond strengthening, whereas in knockout cells, the weaker cytoskeletal network allows stress fibers to slide and concentrate normal forces on individual adhesions, accelerating their rupture.

To validate the model beyond the data used for fitting, we examined its predictions for cell shape under load. The model predicted that synaptopodin-deficient cells would undergo progressive elongation as perpendicular force increased, because weakened actin cables permit asymmetric focal adhesion failure and consequent shape distortion. In contrast, WT cells were predicted to maintain a stable aspect ratio due to uniform force distribution across reinforced adhesion sites. These predictions matched experimental observations: WT cells maintained their aspect ratio of under load, while *Synpo*^⁻/⁻^ cells preferentially lost focal contacts along the thinner dimension of the cell so that the aspect ratio increased (Fig. 5b,d). This was consistent with the rose plot distributions measured experimentally (Fig. 1f-i, Fig. 2f-i), supporting the hypothesis that synaptopodin stabilizes adhesions against perpendicular force.

Mechanistically, in WT cells (Fig. 5d), the high force conversion (*C*) bolsters resistance to perpendicular force around the cell periphery, while *Synpo*^⁻/⁻^ cells lack this force conversion (Fig. 4d, bottom). The individual adhesions with the highest ratio of perpendicular to shear force fail in the absence of force conversion, causing the adhesions closest to the nucleus to detach.

The model thus provides a physical explanation for synaptopodin’s role in perpendicular force resistance. By reinforcing actin bundles, synaptopodin prevents stress fibers from sliding together under normal loading. This maintains geometric separation of focal adhesions and enables force transmission that engages lateral shear-strengthening mechanisms inherent to integrin-based adhesions. Without synaptopodin, perpendicular forces concentrate at individual adhesion sites in a purely normal orientation, leading to rapid bond failure and cell detachment.

## Discussion

Our findings reveal a previously unrecognized form of cellular mechanoadaptation, in which adhesion architecture responds to the direction of applied force. Synaptopodin converts integrin-based adhesions from structures optimized for lateral shear resistance into structures capable of withstanding substantial perpendicular forces through two coordinated mechanisms: reinforcement of actin cables and force-dependent redistribution of adhesion complexes.

The computational model provides insight into the physical basis of this adaptation. A single parameter, *C*, representing efficiency of normal-to-shear force conversion at focal adhesions, distinguishes WT from synaptopodin-deficient cells. We propose that synaptopodin-reinforced actin bundles resist lateral displacement under perpendicular loading, maintaining geometric separation of focal adhesions and distributing forces across multiple attachment sites. This converts a portion of the perpendicular load into lateral shear stress, the force orientation that engages catch-bond strengthening intrinsic to integrin-ligand interactions. Without synaptopodin, actin cables slide under load, causing focal adhesions to concentrate normal forces at individual sites and accelerating bond rupture. The model’s predictive validity extends beyond the fitted detachment data: cell shape changes under load were predicted without additional parameter adjustment, strengthening confidence that the model captures essential physics of synaptopodin-mediated mechanoadaptation.

The peripheral redistribution of β1-integrins represents a striking example of directional mechanosensing, occurring specifically in synaptopodin-expressing cells and only under perpendicular loading. That this reorganization fails entirely in synaptopodin-deficient cells suggests that synaptopodin is required for the mechanosensing pathway itself, not merely for executing the response. Conservation of this mechanism in U2OS cells, which express synaptopodin despite lacking obvious physiological exposure to perpendicular forces, argues that directional mechanoadaptation represents a fundamental cellular capability rather than a podocyte-specific specialization.

The physiological importance of this mechanism is demonstrated by our finding that *Synpo*^⁻/⁻^ mice develop proteinuria, podocyte loss, and foot process broadening when challenged with elevated blood pressure, which increases perpendicular filtrate flow forces. The observation that synaptopodin deficiency produces no phenotype under baseline conditions yet causes podocyte dysfunction under mechanical stress has important clinical implications. This pattern of stress-revealed vulnerability is consistent with previous findings that lack of synaptopodin exacerbates disease in both Adriamycin-induced nephropathy ^11^ and Alport syndrome ^16^, two mechanistically distinct forms of glomerular injury. Together, these studies suggest that deficiencies in mechanoadaptive proteins may create “mechanical vulnerability” in which cells function normally but lack reserve capacity to withstand increased load. Given that synaptopodin downregulation correlates with disease progression in multiple forms of glomerular pathology, therapeutic reinforcement of synaptopodin in injured podocytes could represent a strategy to prevent or slow podocyte detachment.

The sarcomere-like organization of synaptopodin and myosin IIA, and the distinctive structures wrapping around actin filaments in foot processes in vivo, suggest that synaptopodin organizes force-transmitting structures with precise molecular architecture. The colocalization of synaptopodin with β1-integrin at adhesion termini, spatially separated from myosin IIA, implies a division of labor resembling the force-generating and force-transmitting modules of muscle sarcomeres.

Important questions remain, including whether synaptopodin is sufficient to confer perpendicular force resistance to naive cells, and what signaling pathways mediate directional mechanosensing. Nevertheless, our identification of synaptopodin as a molecular switch enabling directional mechanoadaptation reveals a new dimension of cellular mechanobiology with implications for understanding how specialized cells maintain function under diverse mechanical environments. These mechanobiology principles are applicable to numerous clinical settings, including sustained hypertension as a mechanism for accelerated podocyte detachment leading to proteinuria and progression of kidney disease. High intraglomerular pressure states, such as hyperfiltration in diabetes, obesity, sickle cell disease, or low nephron endowment in children, often show early signs of glomerular proteinuria before eventual development of glomerulosclerosis. This is highly relevant, as adequate blood pressure control with proteinuria reduction is the hallmark of therapy to slow progression of chronic kidney disease. Genetic glomerular diseases, such as Alport syndrome and nephrotic podocytopathies, are also relevant to explore, as the sustained high levels or proteinuria are a harbinger of worse kidney outcomes. The onset and worsening of proteinuria might be attenuated therapeutically by taking advantage of our characterization of synaptopodin’s role in mechanoadaptation in podocytes. Our mechanical model provides the groundwork for exploring clinical applications of bolstering synaptopodin and perhaps other podocyte cytoskeletal components in future studies.

## Conclusion

This work establishes directional mechanoadaptation as a fundamental cellular capability and identifies synaptopodin as its molecular basis. By reinforcing actin cables and enabling force-dependent redistribution of adhesion complexes, synaptopodin converts integrin-based adhesions from shear-resistant to perpendicular force-resistant structures, a transformation essential for podocyte survival under glomerular filtration pressures and conserved across diverse cell types. The concept of mechanical vulnerability, in which cells lacking mechanoadaptive proteins function normally at baseline but fail under stress, provides a target for understanding progressive tissue dysfunction in disease. Results demonstrate that cells do actively adapt their structural organization to the direction of applied load, an unexplored dimension of mechanobiology with implications from basic cell physiology to therapeutic strategies for mechanically stressed tissues.

## Acknowledgments

We thank Jennifer Richardson for mouse genotyping and the Genome Engineering & Stem Cell Center for generating the *SYNPO^-/-^* U2OS cells with support from the Washington University Institute of Clinical and Translational Sciences funded by the NIH/National Center for Advancing Translational Sciences, CTSA grant UL1TR002345. This work was funded in part by the NSF through the NSF Science and Technology Center for Engineering Mechanobiology (CMMI 1548571), and by the NIH through grants R01DK131177 (to HYS) and R01DK141178 (to JHM and GMG).

## Methods and Materials

### Fabrication of mechanically tunable hydrogel substrates

Hydroxy-polyacrylamide (PAAm) hydrogels were synthesized using a modified version of previously established protocols.^18^ The main hydrogel solution was prepared by combining HEPES buffer (73.4% v/v, pH 7.4), 40% acrylamide stock solution (16.75% v/v), and 2% bis-acrylamide stock solution (8.0% v/v). Solution was degassed under vacuum for 30 min to remove oxygen that could interfere with polymerization. To initiate crosslinking, we added N-hydroxyethyl acrylamide (HEA, 1.0% v/v), ammonium persulfate (APS, 0.5% v/v), and N,N,N’,N’-tetramethylethylenediamine (TEMED, 0.05% v/v).

Uniform thin hydrogel layers were prepared by sandwiching the reaction mixture between two differentially treated coverslips. Bottom coverslips (22 × 22 mm; Fisher Scientific, 12540B) were cleaned twice with 0.1 M NaOH for 5 min each, rinsed with double-distilled water (ddH₂O), dried, and activated by incubation with 3-(trimethoxysilyl) propyl acrylate for 60 minutes at room temperature. Following activation, coverslips were rinsed with ddH₂O, dried under nitrogen, and either used immediately or stored in a desiccator. Top coverslips (18 × 18 mm) were cleaned with 50% ethanol, rinsed with ddH₂O, dried under nitrogen, and treated with a plasma cleaner (Harrick Plasma PDC-32G) to create a hydrophilic surface. Due to the transient nature of plasma treatment, these coverslips were used within 1 hour.

For polymerization, 30 μL of the complete reaction mixture was placed on an activated bottom coverslip and immediately covered with a plasma-treated top coverslip. After 20 minutes of polymerization, the glass-hydrogel sandwich was transferred to a 6-well plate and submerged in phosphate-buffered saline (PBS) or Hank’s balanced salt solution (HBSS) to quench the reaction. Completed hydrogels were stored at 4°C until use.

### Surface functionalization of hydrogels with laminin-521

Hydrogel surfaces were functionalized with laminin-521, a major GBM component, to promote cell adhesion. The top coverslip was removed from the polymerized hydrogel construct using a razor blade, leaving the hydrogel attached to the activated bottom coverslip. The hydrogel was transferred to a new 6-well plate, washed with sterile HBSS, and sterilized by UV exposure for 60 minutes.

For laminin coating, human laminin-521 (Biolamina, LN521-05) was diluted to 50 μg/mL in PBS containing calcium and magnesium (0.9 mM Ca²⁺, 0.5 mM Mg²⁺), which are required for laminin polymerization and integrin binding. This solution was applied to the hydrogel surface and incubated for 1 hour at room temperature to allow protein adsorption. Excess laminin was removed by gentle rinsing, and the coated hydrogels were dried under a gentle stream of nitrogen gas. Functionalized hydrogels were stored in HBSS-filled 6-well plates at 4°C and used within two weeks.

### Isolation and culture of primary mouse podocytes

All animal experiments were approved by the Washington University in St. Louis Institutional Animal Care and Use Committee (protocol #24-0095). Glomeruli were isolated using magnetic bead perfusion as previously described.^20^ Briefly, 9-month-old WT and *Synpo*^⁻/⁻^ mice on a mixed C57BL/6J - CBA/J background were perfused through the heart with magnetic Dynabeads (150 μL M-450 Epoxy beads, Invitrogen, in 8 mL PBS). Kidneys were harvested, minced, and digested with collagenase A (1 mg/mL; Roche #10103586001) for 30 minutes at 37°C. Digestion was terminated by adding an equal volume of DMEM containing 10% FBS. The tissue suspension was filtered twice through 100-μm strainers (MIDSCI, 220926-299-A) and centrifuged at 290 × *g* (*g* = 9.81 m/s^2^) for 5 min. The pellet was resuspended in HBSS and transferred to a polystyrene tube (Falcon #352052). Bead-containing glomeruli were isolated using a magnetic separation system (EasySep #18000) and collected by washing the tube walls with 1 mL HBSS. Purified glomeruli were cultured in primary podocyte medium for 3 days to allow podocyte outgrowth.

Primary podocyte culture medium was prepared according to established protocols ^22^ by combining: 300 mL conditioned medium from 3T3-L1 cells (collected after 3 days of culture in DMEM with 10% FBS and 1% penicillin-streptomycin), 204 mL low-glucose DMEM (Gibco, 11885084), 102 mL Ham’s F-12 with L-glutamine (Lonza #12-615F), 30 mL FBS, 6 mL penicillin-streptomycin (Sigma-Aldrich #P4333), and 6 mL insulin-transferrin-selenium supplement (Invitrogen #41400045).

To isolate podocytes from outgrowth cultures, medium was removed and cultures were washed once with HBSS, then incubated with Cell Stripper (Corning #25-056-CL) for 30 minutes at 37°C. Cells were detached by gentle pipetting and filtered through a 40-μm strainer (MIDSCI, 230304-301-A) to exclude glomerular cores. Podocytes in the filtrate were collected by centrifugation and resuspended in fresh primary podocyte medium.

### Application of perpendicular and lateral shear forces to cultured cells

For perpendicular force experiments, custom sample holders were designed using biocompatible resin (BioMed Clear V1) and fabricated on a Formlabs (Somerville, MA) Form 3B+ 3D printer (Supplementary Fig. 1a). Laminin-coated hydrogel constructs were mounted in these holders and placed in 60 mm tissue culture dishes (TPP #93060). Dishes were filled with pre-warmed HBSS (37°C), sealed with parafilm, enclosed in plastic bags, and mounted in pairs in a temperature-controlled swinging-bucket centrifuge (Eppendorf 5810 R) maintained at 37°C. Centripetal accelerations ranging from 30*g* to 450*g* (*g* = 9.81 m/s^2^) were applied by adjusting centrifugation speed. Net normal stress on cells was estimated as:

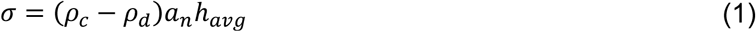

where *ρ*_*c*_ is the density of cells (∼1.05 g/mL), *ρ*_*d*_ is the density of the medium (∼1.01 g/mL), *α*_*n*_ is the applied centripetal acceleration, and ℎ_*avg*_ is the average cell height. This calculation accounts for the buoyancy of cells in the surrounding medium.

For lateral shear force experiments, microfluidic chambers with defined channel dimensions were fabricated using the same 3D printing approach (Supplementary Fig. 1b). Hydrogel constructs were mounted in chamber slots, and controlled laminar flow was applied using an InfusionONE syringe pump (New Era Pump Systems #98242) equipped with a 100 mL syringe. Wall shear stress (*τ*) was calculated from the flow rate and channel cross-sectional geometry, and adjusted by modulating flow rate.

### Cell migration assay

Cell migration was assessed using a wound closure assay. WT and *SYNPO*^⁻/⁻^ U2OS cells were seeded into removable two-well culture inserts (Ibidi, 81176) and cultured overnight in low-serum medium (DMEM supplemented with 0.1% FBS and 1% penicillin-streptomycin) to minimize the contribution of cell proliferation to gap closure. Initial cell confluence was approximately 90% immediately after seeding and reached 100% after overnight culture. Once cells reached full confluence, the silicone insert was removed to create a defined cell-free gap, and cells were allowed to migrate into the gap. Phase-contrast images were acquired at 0, 3, 6, 9, 12, 15, 18, 21, and 24 hours post-insert removal to monitor gap closure over time.

### Immunofluorescence microscopy

Following force application, cells on hydrogels were fixed with 4% paraformaldehyde in PBS (Electron Microscopy Sciences #15712) for 10 minutes at room temperature. Samples were washed three times with PBS (5 minutes each), permeabilized with 0.05% Triton X-100 in PBS for 10 minutes, and blocked with 2% bovine serum albumin (BSA; Sigma-Aldrich #A7906) in PBS for 30 minutes. Primary antibodies diluted in blocking buffer were applied overnight at 4°C.

After three PBS washes, samples were incubated with species-appropriate fluorophore-conjugated secondary antibodies for 1 hour at room temperature. Following three final PBS washes, samples were mounted in SlowFade antifade medium (Invitrogen #S36917) and sealed with nail polish.

The following primary antibodies were used: anti-synaptopodin (Synaptic Systems #163004 for mouse samples; American Research Products #03-GP94-N, N-terminal-specific, for mouse and human samples), anti-myosin IIA (BioLegend #909801), anti-α-actinin-4 (Abcam #ab68167), anti-β1-integrin (BD Pharmingen #553715; mouse-specific), anti-podocalyxin (R&D Systems #AF1556), and anti-WT1 (Abcam #ab224806). F-actin was visualized using Alexa Fluor 647-conjugated phalloidin (Thermo Fisher #A22287). Secondary antibodies were selected to match primary antibody host species and conjugated to spectrally compatible fluorophores for multicolor imaging.

### Image acquisition and analysis

Images were acquired using a Nikon spinning disk confocal microscope equipped with a Yokogawa CSU-X1 scan head, LED-based DMD illumination system, and Andor Zyla sCMOS camera. Cell detachment assays were imaged with a 10× objective (NA 0.45) to capture large fields of view, whereas protein localization studies used a 60× oil-immersion objective (NA 1.4) for high-resolution imaging. All imaging parameters, including exposure time, laser power, and detector gain, were kept constant within experimental groups to enable quantitative comparisons.

Image analysis was performed using ImageJ (NIH). For cell detachment assays, images were processed by background subtraction and threshold adjustment, followed by automated cell counting. Cell density was calculated as the number of cells per unit area.

For protein localization studies, focal adhesion distributions were quantified using a custom peripheral localization index that measured the relative concentration of β1-integrin signal at the cell periphery compared to the cell center. For each cell, a line profile of fluorescence intensity was drawn from one edge of the cell through the centroid to the opposite edge. The intensity profile *I*(*x*) was normalized over a standardized length (1000 points) using min-max normalization:

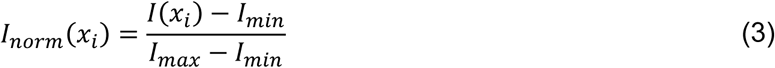

where *I*(*x*_*i*_) is the raw fluorescence intensity at point *i* along the profile, and *I_min_* and *I_max_* are the minimum and maximum intensity values along the entire profile.

To quantify peripheral versus central distribution, we calculated the mean intensity in the peripheral regions (the outer 10% at each end of the profile) relative to the central region (the middle 10% of the profile):

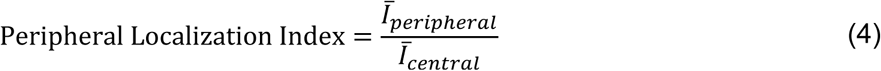

where:

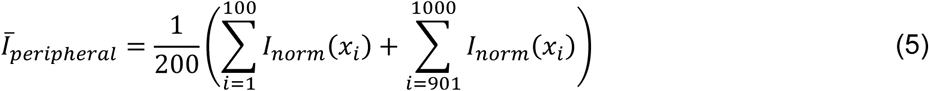

And

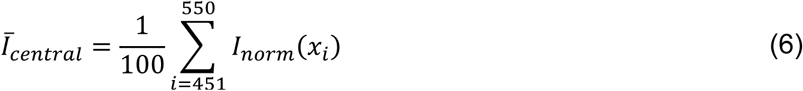

A peripheral localization index greater than 1 indicates enrichment of focal adhesions at the cell periphery, while a value less than 1 indicates central concentration. For each experimental condition, at least 30 cells were analyzed, and line profiles were drawn along the major axis of elongated cells or along a random diameter for rounded cells.

### Protein immunoblot analysis

WT and *SYNPO^-/-^* U2OS cells were cultured to 90% confluence in 100 mm tissue culture dishes (TPP). Cells were washed once with cold PBS, then lysed directly in 2× Laemmli sample buffer (Bio-Rad #1610747) supplemented with 10% 2-mercaptoethanol. Lysates were collected by scraping, transferred to microcentrifuge tubes, and denatured at 95°C for 5 minutes. Proteins were resolved on 4–20% gradient Criterion TGX precast gels (Bio-Rad #5671095) by SDS-PAGE at 125 V for 1 hour, then transferred to nitrocellulose membranes (Bio-Rad #1620112) using a wet transfer system at 40 V for 2 hours at 4°C.

Membranes were blocked with 5% non-fat dry milk in PBST (PBS containing 0.2% Tween-20; Fisher BioReagents, BP337-500) for 30 minutes at room temperature with gentle agitation. Primary antibodies diluted in blocking buffer were applied overnight at 4°C. The following primary antibodies were used: anti-synaptopodin (American Research Products #03-GP94-N), anti-α-actinin-4 (Abcam #ab68167), and anti-GAPDH (Cell Signaling Technology #2118) as a loading control. After three 10-minute washes in PBST, membranes were incubated with species-appropriate horseradish peroxidase-conjugated secondary antibodies for 1 hour at room temperature. Following three additional PBST washes, immunoreactive bands were detected by enhanced chemiluminescence using a ChemiDoc MP imaging system (Bio-Rad #12003154). Densitometric quantification of band intensities was performed using ImageJ (NIH), with protein levels normalized to GAPDH.

### AAV-mediated hypertension model and assessment of kidney function

Adeno-associated virus serotype 8 vectors expressing renin (AAV8-TBG-Ren1) or GFP (AAV8-TBG-GFP) under the hepatocyte-specific thyroxine-binding globulin (TBG) promoter were produced by the Hope Center Viral Vectors Core at Washington University School of Medicine. Viruses were diluted in sterile PBS to a final dose of 5 × 10^10^ viral genomes per mouse and administered by retro-orbital injection under isoflurane anesthesia.

Urine samples were collected immediately before injection (week 0) and at 2, 4, and 6 weeks post-injection. Urinary creatinine concentration was measured using a colorimetric assay (BioAssay System, #DICT-500), and urinary albumin was quantified by SDS-PAGE alongside albumin standards of known concentration. Albumin-to-creatinine ratios were calculated for each sample. At week 6, mice were euthanized and kidneys were harvested for electron microscopy and immunofluorescence analysis. Livers were collected to verify renin expression by immunostaining.

Arterial blood pressure was measured under 1.5% isoflurane anesthesia as previously described using a Millar catheter (model SPR-1000) introduced into the ascending aorta via the right common carotid artery. ^27^

### Quantification of podocyte number *in vivo*

Kidney tissue was embedded in Tissue-Tek O.C.T. compound (Sakura #4583), frozen, and sectioned at 7 μm using a cryostat (Leica CM1950). Cryosections were immunostained with anti-WT1 to identify podocyte nuclei and anti-podocalyxin to delineate glomerular boundaries (Supplementary Fig. 2a-d).

Podocyte density was quantified using a combination of ImageJ (NIH) and MATLAB (R2020b, MathWorks, Natick, MA). Glomerular boundaries were manually traced in ImageJ based on podocalyxin staining, and the enclosed regions were converted to binary masks for area calculation. WT1-positive nuclei within each glomerular boundary were counted manually. Podocyte density was calculated as the number of WT1-positive nuclei divided by glomerular cross-sectional area (Supplementary Fig. 2e, f). At least 20 glomeruli were quantified per mouse.

### Electroporation of synaptopodin long isoform

*Synpo*^⁻/⁻^ primary podocytes were isolated as described above. Electroporation buffer was prepared according to established protocols,^21,26^ consisting of 5 mM KCl, 15 mM MgCl_2_, 120 mM Na_2_HPO_4_, and 50 mM mannitol. For electroporation, podocytes were resuspended in 200 μL of transfection mixture containing 166 μL electroporation buffer, 20 μL of 1 M NaCl, 4 μL of 10% poloxamer 188 (Sigma-Aldrich, P5566), and 10 μL of plasmid DNA (5 μg total) encoding the synaptopodin long isoform.

Cells were transferred to a 4 mm electroporation cuvette (Bio-Rad, 1652088) and electroporated using a Gene Pulser Xcell system (Bio-Rad) with the following parameters:^23^ 140 V, square wave pulse, 25 ms duration. Following electroporation, cells were immediately transferred to pre-warmed primary podocyte medium and plated on laminin-521-coated hydrogels for recovery and subsequent analysis.

### Nanofountain probe electroporation of synaptopodin long isoform

Single-cell electroporation was performed using the nanofountain probe electroporation (NFP-E) system as previously described.^18,19,24,25^ *Synpo^-/-^* primary podocytes were isolated and cultured on laminin-521-coated hydrogels as described above. After 48-72 hours of culture, cells were transferred to the NFP-E platform mounted on an inverted fluorescence microscope (Nikon Eclipse Ti-E) for transfection.

Glass nanopipettes (Eppendorf) with tip apertures of approximately 100-200 nm were back-filled with plasmid DNA encoding the synaptopodin long isoform (0.5-1 μg/μL in 1X PBS). The nanopipette was mounted on a robotic arm controlled by precision XYZ piezo stages (40 nm resolution). Individual cells were identified using phase contrast microscopy, and the nanopipette was positioned over selected cells using either manual selection or automated cell localization via a deep convolutional neural network.^19, 25^

Cell-nanopipette contact was established using an automated resistance-based detection algorithm. As the nanopipette approached the cell membrane, system resistance was continuously monitored at 150 Hz. Contact was confirmed when the resistance increased above a threshold of 1% of baseline (typically 300-500 kΩ above baseline resistance of 30-50 MΩ).

Upon contact detection, bilevel electroporation pulses were applied (first level: 15 V, 0.5 ms; second level: 10 V, 2.5 ms) at a frequency of 50 Hz. Two pulse trains of 50 pulses each were delivered with a 1-second interval between trains. Following electroporation, the nanopipette was retracted and repositioned to the next target cell.

This single-cell approach enabled mosaic transfection, where only selected cells within the culture received plasmid DNA while neighboring cells remained untransfected, serving as internal controls. After electroporation, cells were returned to primary podocyte culture medium and incubated at 37°C with 5% CO_2_ for 24-48 hours before immunofluorescence analysis. Cell viability was assessed by morphological criteria and propidium iodide exclusion.

### Computational model of cell detachment

To gain mechanistic insight into how synaptopodin enables perpendicular force resistance, we developed a computational model of cell detachment under inertial loading. The model integrates three key components: (1) cell geometry and stress fiber architecture, (2) mechanical equilibrium and force transmission through stress fibers, and (3) force-dependent focal adhesion kinetics. A single free parameter, *C*, representing the efficiency of normal-to-shear force conversion at focal adhesions, distinguishes WT from synaptopodin-deficient cells. The model was implemented in MATLAB (R2020b, MathWorks) and validated against experimental measurements of cell detachment and shape change.

*Cell geometry and stress fiber architecture.* The cell was modeled as a mechanical system comprising a rigid nucleus connected to the substrate via discrete stress fibers. The outer cell boundary was defined as an ellipse with major axis *α* and minor axis *b*. For a stress fiber oriented at an angle *θ* relative to the major axis, the radial distance to the cell margin *r*(*θ*)was calculated as:

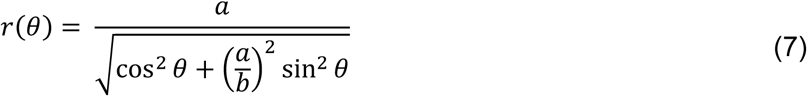

To isolate the effects of cell shape from cell size, the cytoplasmic area was held constant across varying aspect ratios (*AR*). Cell dimensions were scaled such that 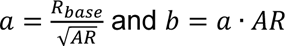, where *R*_*base*_ is the baseline radius. The nucleus was idealized as a sphere with a circular actin cap of radius *r*_0_ positioned at a height ℎ above the substrate. Stress fibers were modeled as elastic cables connecting the actin cap to focal adhesions at the cell periphery. The unstretched length *L*_0_ of a fiber at angle *θ* was defined by the geometric hypotenuse:

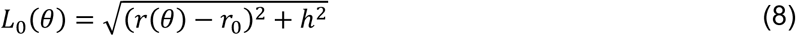

The spatial distribution of stress fibers was determined by a polarization factor to reflect cytoskeletal organization observed experimentally. For elongated cells (*AR* > 1.5), fiber angles were distributed according to a weighted probability density function that biases fibers toward the major axis, simulating anisotropic cytoskeletal organization. Circular cells (*AR* ≤ 1.5) used a uniform random distribution.

### Mechanical equilibrium and force transmission

Each stress fiber *i* was treated as a linear spring with stiffness *k*_*i*_ inversely proportional to its unstretched length:

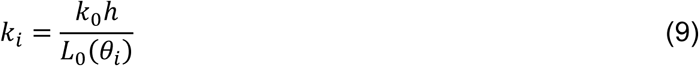

where *k*_0_ is a reference stiffness. Upon application of an inertial force *F*_*inertia*_to the nucleus (simulating centrifugation), the system undergoes a vertical displacement *δ*The stretched length of fiber *i* at time step *j* becomes:

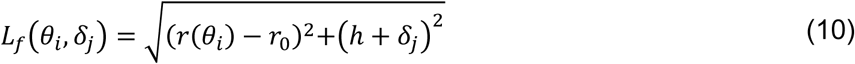

The tension *F*_*i*_ in each fiber was calculated as the sum of a constant actomyosin contractile force *F*_0_ and the passive elastic force resulting from extension:

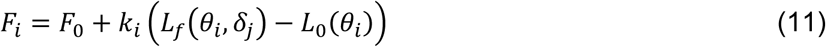

This tension is resolved into a normal component *N*_*i*_ = *F*_*i*_ sin*ϕ*_*i*_ and a lateral shear component *S*_*i*_ = *F*_*i*_ cos *ϕ*_*i*_ at the focal adhesion interface. At every time step, the vertical displacement *δ*_*j*_ was solved iteratively to satisfy mechanical equilibrium, where the sum of the vertical force components of all bonded fibers balances the applied inertial load:

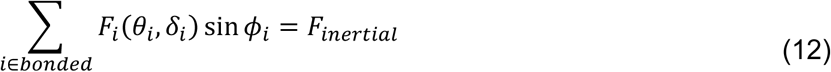

### Focal adhesion kinetics

The stability of focal adhesions was modeled using a force-dependent catch-slip dissociation rate. The off-rate *k_off_* for each bond was modulated by both normal and lateral shear forces according to the following relation:

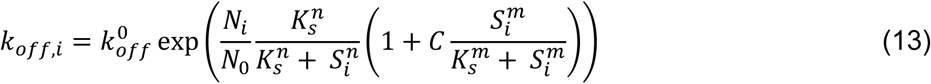

Here, *C* represents a specific molecular parameter (modulated in the simulation to represent different experimental conditions, e.g., Control vs. Knockout), while *N*_0_, *K*_*s*_, *K*_0_, *n*, and *m* are constants governing the bond sensitivity. The probability of a specific focal adhesion debonding within a time step Δ*t* was calculated as *P*_*i*_ = 1 − exp(−*k_off_*,*i*Δ*t*).^10^

### Simulation parameters and protocol

Monte Carlo simulations were performed using MATLAB (The MathWorks, Natick, MA). For each condition, *N* = 500 independent trials were conducted, with each cell containing *N_fibers_* = 50 initially bonded fibers. The simulation swept through force magnitudes (*F*_*inertial*_) ranging from 1 × to 15 × a baseline force (*F*_*base*_ = 5 nN). A cell was considered “detached” if the number of remaining bonds fell below 5% of the initial population (< 0.05 *N_fibers_*) The parameter *C* was varied between 1.3 and 2.45 to simulate different adhesion strengths.

---

**Supplementary Fig. 1.**
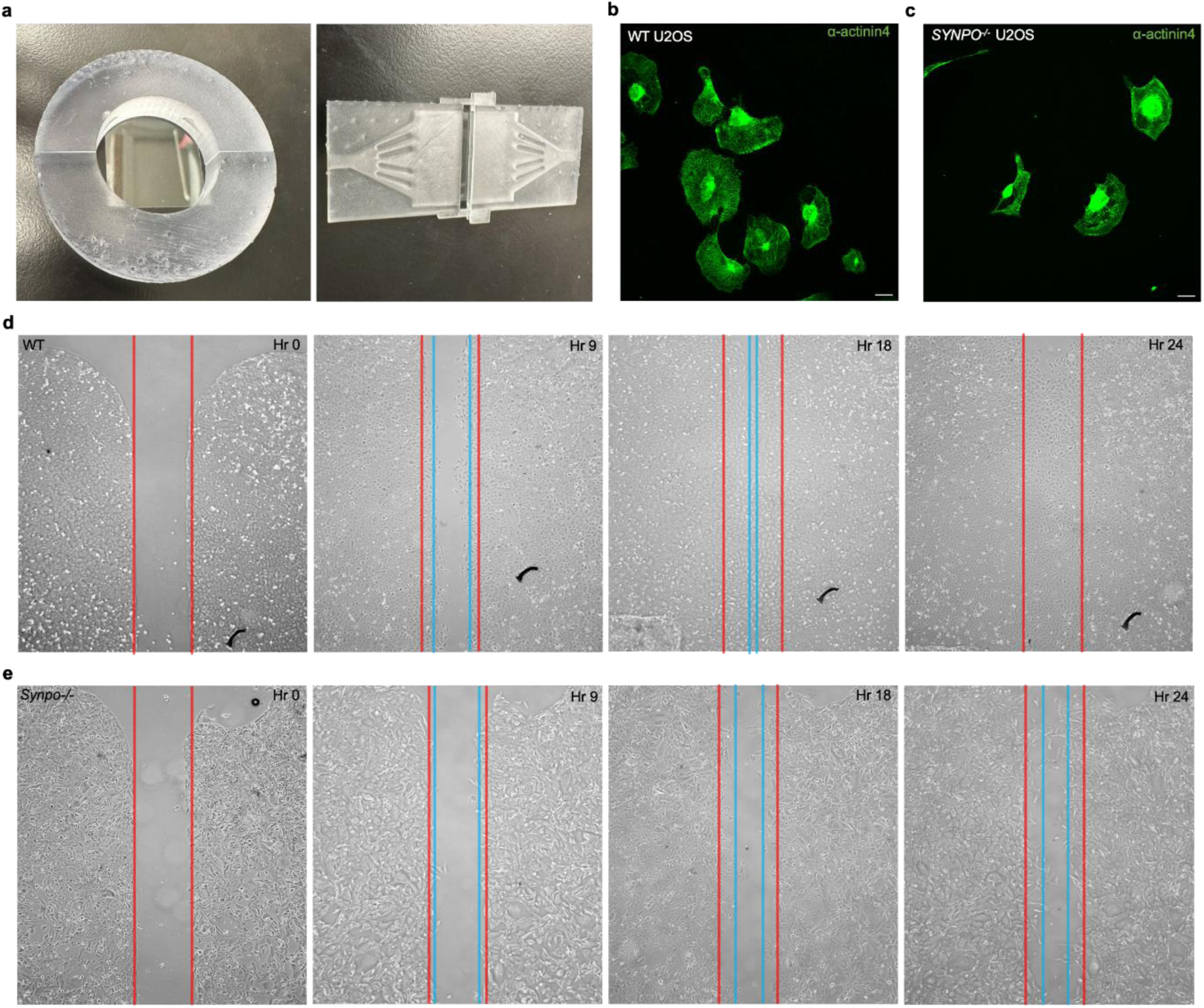
Experimental apparatus for force application and additional characterization of *SYNPO^-/-^* U2OS cell defects. **(a)** Custom-designed sample holders for applying defined mechanical forces to cells cultured on hydrogel substrates. Left: perpendicular force apparatus consisting of 3D-printed holders (BioMed Clear V1 resin) that secure hydrogel constructs in tissue culture dishes for centrifugation in a temperature-controlled swinging bucket rotor. Right: lateral shear force apparatus consisting of a microfluidic chamber with defined channel dimensions that generates controlled laminar flow across the cell surface when connected to a syringe pump. Both systems maintain cells at 37°C in buffered saline throughout force application. **(b, c)** Representative immunofluorescence images of WT **(b)** and *SYNPO*^⁻/⁻^U2OS cells stained for α-actinin-4 (green) following perpendicular force application. WT cells maintain continuous cytoskeletal networks with α-actinin-4 distributed along intact stress fibers. *SYNPO*^⁻/⁻^ cells display pronounced cytoskeletal fractures characterized by gaps in α-actinin-4 staining, particularly in perinuclear regions, consistent with the mechanical vulnerability observed in main Figure 2. **(d, e)** Wound closure migration assay comparing WT **(d)** and *SYNPO*^⁻/⁻^ **(e)** U2OS cells. Cells were cultured to confluence in two-well silicone inserts; insert removal created a defined gap at time *t* = 0. Images were acquired at 0, 9, and 24 hours. WT cells migrated into the gap within 9 hours and achieved complete closure by 24 hours. *SYNPO*^⁻/⁻^ cells showed markedly impaired migration, with minimal gap narrowing at 9 hours and persistent gaps at 24 hours. This motility defect is consistent with compromised adhesion dynamics in synaptopodin-deficient cells. Scale bars: 20 μm **(b, c)**; 20 μm **(d, e)**.

**Supplementary Fig. 2.**
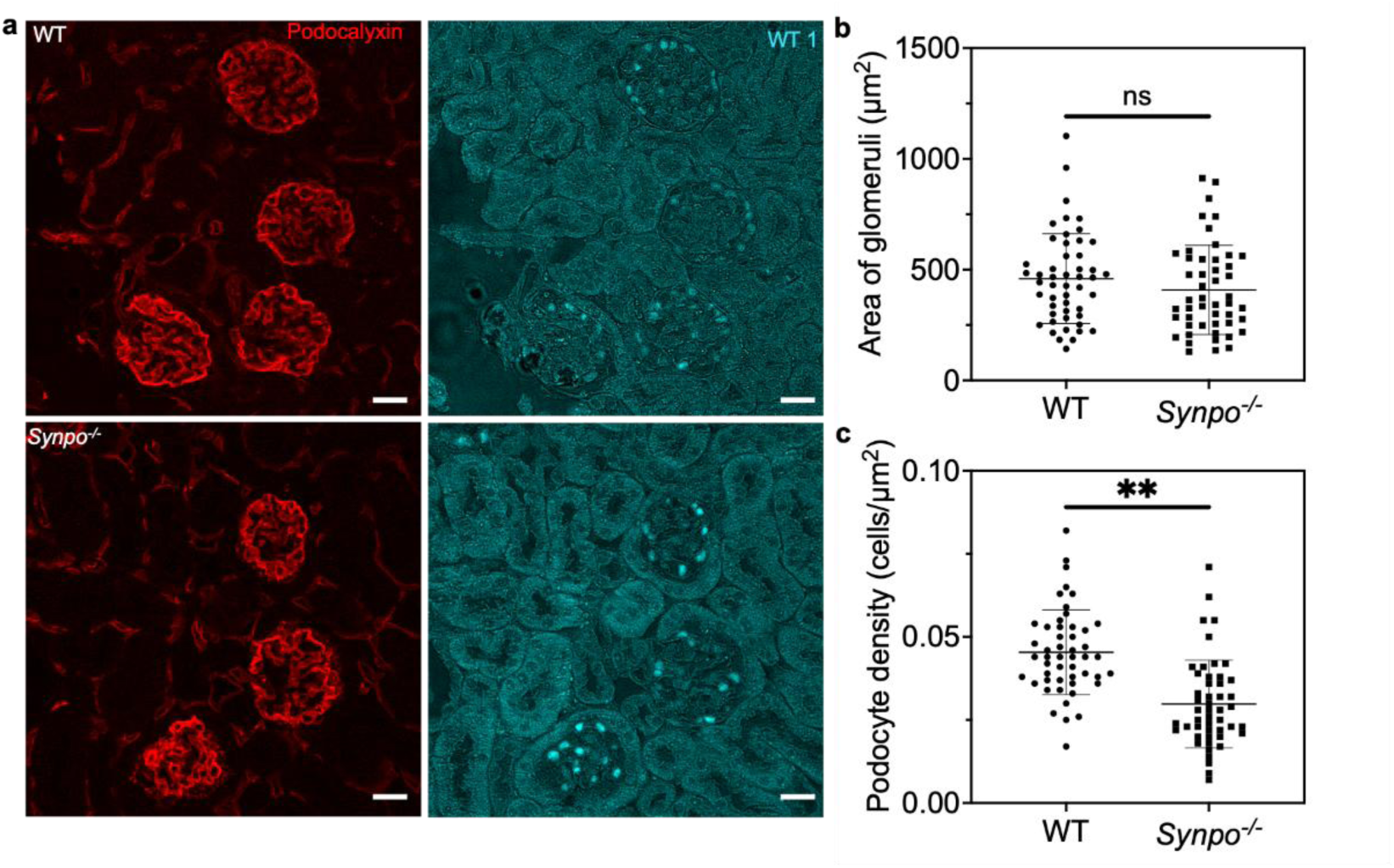
Elevated blood pressure causes podocyte loss in *Synpo^-/-^* but not WT mice. **(a, b)** Representative immunofluorescence images of kidney cryosections from AAV8-Renin-injected WT **(a)** and *Synpo*^⁻/⁻^ **(b)** mice, stained for podocalyxin (green) to delineate glomerular boundaries and WT1 (red) to identify podocyte nuclei. Glomeruli from *Synpo*^⁻/⁻^ mice contain visibly fewer WT1-positive cells compared to WT controls despite similar glomerular size. **(c, d)** Higher magnification views of representative glomeruli from WT **(c)** and *Synpo*^⁻/⁻^ **(d)** mice illustrating the reduction in podocyte number. Dashed lines indicate glomerular boundaries traced for area quantification. **(e)** Quantification of glomerular cross-sectional area. No significant difference was observed between WT and *Synpo*^⁻/⁻^ mice, indicating that podocyte loss is not secondary to glomerular hypoplasia or hypertrophy. **(f)** Quantification of podocyte density (WT1-positive nuclei per glomerular area). *Synpo*^⁻/⁻^ mice exhibit significantly reduced podocyte density compared to WT mice following AAV8-Renin treatment, consistent with podocyte detachment under elevated perpendicular filtration forces. This podocyte loss likely contributes to the proteinuria observed in *Synpo*^⁻/⁻^ mice (Fig. 3b) and parallels the in vitro detachment phenotype (Fig. 1d). Glomerular areas and podocyte numbers were quantified using automated image analysis in MATLAB. Data represent mean ± SEM; *n* ≥ 5 mice per group; at least 20 glomeruli quantified per mouse. Scale bars: 20 μm **(a, b)**; 20 μm **(c, d)**.

**Supplementary Fig. 3.**
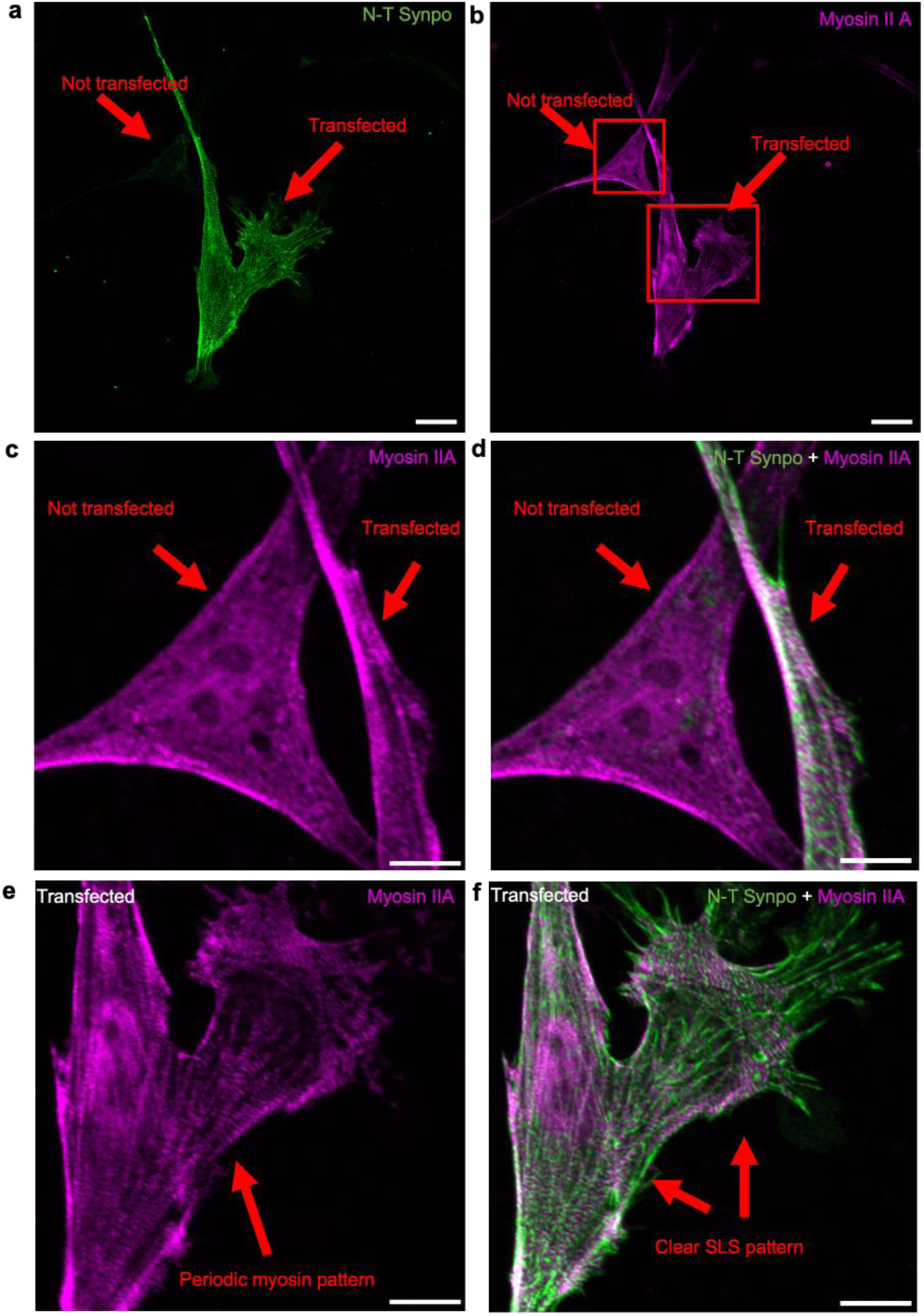
Synaptopodin expression is sufficient to organize myosin IIA into sarcomeric patterns in primary podocytes. **(a, b)** Low-magnification immunofluorescence images of *Synpo*^⁻/⁻^ primary podocytes transfected with the synaptopodin long isoform (Synpo-L) using nanofountain probe electroporation (NFP-E) and stained for synaptopodin using an N-terminal-specific antibody (green) **(a)** and myosin IIA (MyoIIA; purple). **(b)** Mosaic expression allows direct comparison of transfected and non-transfected cells within the same field. **(c, d)** Higher magnification of a non-transfected *Synpo*^⁻/⁻^ podocyte showing MyoIIA alone **(c)** and merged with synaptopodin staining **(d)**. In the absence of synaptopodin, MyoIIA displays diffuse, disorganized distribution with no discernible periodic structure. **(e, f)** Higher magnification of a neighboring transfected podocyte showing MyoIIA alone **(e)** and merged with synaptopodin staining **(f)**. Following synaptopodin expression, MyoIIA adopts a sarcomeric pattern, forming alternating bands with synaptopodin along stress fibers. This cell-autonomous reorganization demonstrates that synaptopodin is sufficient to organize the actomyosin cytoskeleton into the sarcomere-like structures (SLSs) observed in WT cells (Fig. 4b) and suggests that synaptopodin acts as a structural template for contractile machinery assembly. The single-cell transfection approach provides an internal control, as transfected and non-transfected cells experience identical culture conditions and imaging parameters. Scale bars: 20 μm **(a, b)**; 10 μm **(c–f)**.

## Notes

### Competing Interest Statement

The authors have declared no competing interest.

